# Ecm29-dependent proteasome localization regulates cytoskeleton remodeling at the immune synapse

**DOI:** 10.1101/2020.12.23.424144

**Authors:** Jorge Ibañez-Vega, Felipe Del Valle, Juan José Saez, Jheimmy Diaz, Andrea Soza, María Isabel Yuseff

**Author notes:** These authors contributed equally to this work (Jorge Ibañez-Vega, Felipe Del Valle). **Correspondence:** Andrea Soza, María Isabel Yuseff.

## Abstract

The formation of an immune synapse (IS) enables B cells to capture membrane-tethered antigens, where cortical actin cytoskeleton remodeling regulates cell spreading and depletion of F-actin at the centrosome promotes the recruitment of lysosomes to facilitate antigen extraction. How B cells regulate both pools of actin, remains poorly understood. We report here that decreased F-actin at the centrosome and IS relies on the distribution of the proteasome, regulated by Ecm29. Silencing Ecm29 decreases the proteasome pool associated to the centrosome of B cells and shifts its accumulation to the cell cortex and IS. Accordingly, Ecm29-silenced B cells display increased F-actin at the centrosome, impaired centrosome and lysosome repositioning to the IS and defective antigen extraction and presentation. Ecm29-silenced B cells, which accumulate higher levels of proteasome at the cell cortex, display decreased actin retrograde flow in lamellipodia and enhanced spreading responses. Our findings support a model where B the asymmetric distribution of the proteasome, mediated by Ecm29, coordinates actin dynamics at the centrosome and the IS, promoting lysosome recruitment and cell spreading.

## Introduction

The interaction of the B cell receptor (BCR) with membrane-tethered antigens (mAg) initiates the formation of an immune synapse (IS), characterized by rapid cortical actin cytoskeleton rearrangements and the formation of BCR-microclusters containing signaling molecules that elicit B cell activation (Yuseff *et al*., 2013; Kwak, Akkaya and Pierce, 2019). In resting conditions, the lateral diffusion of the BCR is restricted by the cortical actin network, which becomes disassembled upon antigen engagement, enabling BCR lateral diffusion and subsequent clustering to promote downstream signaling (Freeman et al., 2018; Mattila, Batista, & Treanor, 2016; Tolar, 2017). During this phase, B cells exert a rapid spreading response by forming lamellipodia to expand the contact area with the antigen-presenting surface, thereby increasing the number of BCRs coupled to mAg (Spillane & Tolar, 2018). IS formation also involves the mobilization of the centrosome together with lysosomes towards the Ag-contact site, where secretion of their content into the synaptic space facilitates antigen extraction from stiffer surfaces (Yuseff *et al*., 2011). Translocation of the centrosome from the perinuclear region to the IS relies on actin remodeling, where depolymerization of the actin pool surrounding the centrosome allows its uncoupling from the nucleus, thereby promoting the establishment of a polarized phenotype (Obino *et al*., 2016). Thus, the capacity of B cells to organize an IS and execute their antigen-presenting function is strongly dependent on actin remodeling both at the cortical and perinuclear regions. How B cells orchestrate actin dynamics at both these levels remains to be resolved.

The ubiquitin-proteasome system (UPS) has emerged as a critical regulator of cell signaling, polarization, cell division, and migration by selective proteolysis of ubiquitin-tagged proteins (Coux, 2002; Schaefer, Nethe and Hordijk, 2012; Collins and Goldberg, 2017). This system comprises ubiquitin ligases that targets proteins for degradation by covalently conjugating them with ubiquitin, enabling recognition by the proteasome to drive their proteolysis (Collins and Goldberg, 2017). The proteasome is responsible for the degradation of most cytosolic proteins in mammalian cells. This protein complex is formed by the 20S core particle (CP) and the 19S regulatory particle (RP), that caps the 20S CP on one (26S proteasome) or both ends (30S proteasome) in an ATP-dependent manner and can dissociate reversibly (Collins and Goldberg, 2017). Proteasome assembly, activity, localization, and half-life is regulated by transcriptional and post-translational modifications of proteasomal subunits (Dahlmann, 2016; Collins and Goldberg, 2017; Dikic, 2017).

Among proteasome regulators, a 200 kDa protein, termed Ecm29, first characterized in yeast, binds the proteasome to motor proteins and vesicles, suggesting that it could play a role in the intracellular localization of the proteasome. Ecm29 has been shown to couple the proteasome to the Endoplasmic Reticulum (ER), microtubules, and centrosome (Gorbea *et al*., 2010), however, the mechanisms by which Ecm29 recruits the proteasome to specific cellular compartments are not fully understood. In neurons, Ecm29 controls proteasome localization and mobilization across the axon by modulating its association with microtubules, where its activity and localization influences neuronal development and synaptic signaling (Otero *et al*., 2014; Hsu *et al*., 2015; Pinto *et al*., 2016; Lee *et al*., 2020).

In lymphocytes, proteasome activity and localization also play a crucial role in their function. Indeed, the inhibition of proteasome activity leads to defective actin remodeling and reduced ERK signaling, impairing efficient B and T lymphocyte activation (Schmidt *et al*., 2018; Ibañez-Vega, Del Valle Batalla, *et al*., 2019). Moreover, during asymmetric T cell division, the unequal segregation of the proteasome between the two daughter cells enables the selective degradation of the transcriptional factor Tbet, which ultimately leads to the acquisition of different phenotypes (Dennis *et al*., 2012). Thus, the localization of the proteasome is regulated in lymphocytes during asymmetric cell division, where control of cell polarity is critical. Analogously, in neurons, where cell polarity is also crucial, the localization of the proteasome targets the degradation of ubiquitylated proteins, required for axon development (Otero *et al*., 2014; Hsu *et al*., 2015) and presynaptic differentiation (Pinto *et al*., 2016; Liu *et al*., 2019). Here, a polarized phenotype is achieved by the selective degradation of polarity proteins, such as PAR-2, PAR-3, PAR-6 (Laumonnerie and Solecki, 2018), and actin polymerizing factors, such as VASP by the UPS (Boyer *et al*., 2020).

Interestingly, upon activation, B cells upregulate the ubiquitylation of proteins, including BCR downstream signaling molecules, polarity proteins, and actin polymerizing factors (Satpathy *et al*., 2015), highlighting a role for the UPS in regulating actin dynamics and B cell activation. We have previously shown that B cells contain an active proteasome pool at the centrosome, which is required for efficient actin clearance at this level, which enables centrosome repositioning to the immune synapse (Ibañez-Vega, Del Valle Batalla, *et al*., 2019). However, the underlying mechanisms of proteasome localization remain to be addressed in lymphocytes.

In this study, we explored whether Ecm29 controls the localization of the proteasome in B cells during the formation of an immune synapse and how this specific localization coordinates actin remodeling responses between the synaptic interface and the centrosome. Our results show that Ecm29 mediates the association of the proteasome with the microtubule network and regulates the distribution of proteasome pools at the centrosome and the immune synapse of B cells. Upon Ecm29 silencing, B cells redistribute the proteasome to the synaptic membrane, which results in defective actin dynamics at this level, evidenced by slower actin retrograde flow at lamellipodia formed at the synaptic membrane and increased spreading responses. Ecm29-silenced B cells also displayed deficient actin depolymerization at the centrosome, which impaired centrosome and lysosome repositioning at the immune synapse, resulting in reduced antigen extraction and presentation. Overall, our results contribute to understanding how B lymphocytes efficiently coordinate actin dynamics at the centrosome and the synaptic interface by controlling the localization of the proteasome. We propose that the distribution of the proteasome depends on Ecm29, which enables: 1-the accumulation of the proteasome at the centrosome, used to promote actin depolymerization required for centrosome re-positioning and lysosome recruitment at the IS. 2-the recruitment of proteasome to the synaptic membrane, promoting actin depolymerization at this level to enhance cell spreading and signaling.

Thus, Ecm29 emerges as a key regulator of proteasome distribution used to orchestrate key synaptic functions to facilitate antigen extraction and activation of B cells.

## Results

### Ecm29 regulates the localization of the proteasome in B cells

Intracellular compartmentalization of proteasome activity controls actin cytoskeleton remodeling in B cells during immune synapse formation (Ibañez-Vega, et al. 2019). We sought for potential regulators involved in proteasome distribution and focused on the molecule Ecm29, a proteasome adaptor, and scaffold protein, which binds to the 26S proteasome and links it to motor proteins, vesicles, centrosome, and microtubules (Gorbea *et al*., 2010). To this end, we first analyzed the localization of Ecm29 in resting and activated B cells using immobilized antigens, which trigger the formation of an IS (Yuseff *et al*., 2013; Ibañez-Vega, Fuentes, *et al*., 2019). We found that upon activation with antigen-coated beads, used to mimic the formation of an IS, Ecm29 progressively accumulated at the synaptic interface (**Figure 1A and B**), similarly to the proteasome (Ibañez-Vega, Del Valle Batalla, *et al*., 2019). To further characterize the distribution of Ecm29 at the synaptic membrane, we activated B cells on antigen-coated coverslips and labeled Ecm29 together with microtubules (α-tubulin) and the centrosome (centrin-GFP). We found that Ecm29 was distributed in a central and peripheral pool, associated with the centrosome and cortical microtubules, respectively (**Figure 1C**). The accumulation of Ecm29 at the centrosome of B cells was verified by immunoblot of centrosome-rich fractions, where we found that Ecm29 cofractionated with γ-tubulin, a centrosome marker (**Figure EV1**). Imaging analysis also revealed that Ecm29 colocalized with microtubules, displaying a Pearson’s coefficient over 0.8 in resting and activating conditions (**Figure 1C and D)**, suggesting that Ecm29 is tightly associated with the microtubule network. Indeed, the association of Ecm29 with microtubules was previously described in neurons, where it was shown to mediate proteasome retrograde transport and axon development (Otero *et al*., 2014; Hsu *et al*., 2015). Interestingly, we found that upon activation, Ecm29 slightly reduced its colocalization with microtubules and accumulated at the cell periphery (**Figure 1C, D, and E**), suggesting that Ecm29 changes its distribution in response to BCR stimulation.

**Figure 1:**
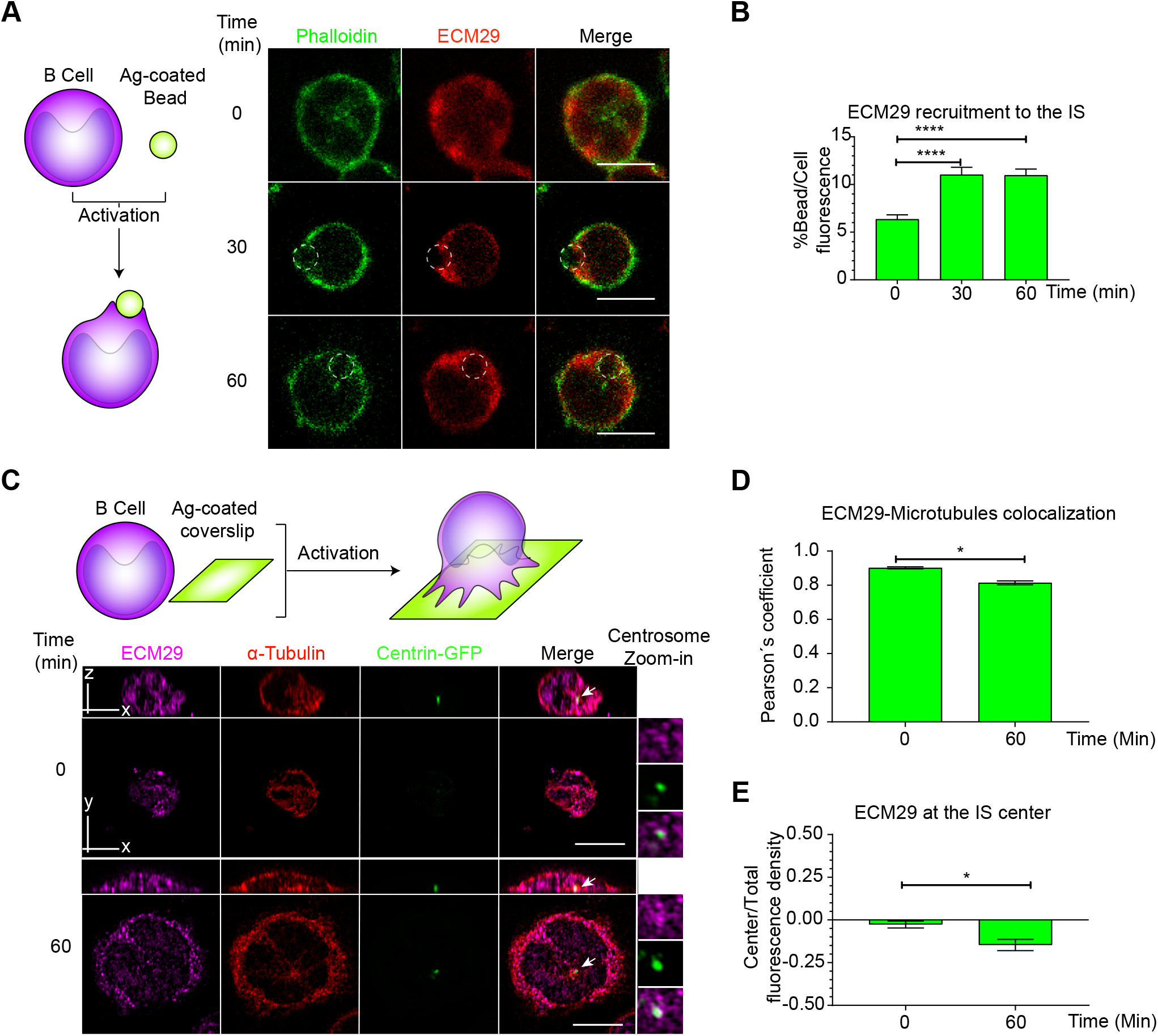
Ecm29 co-localizes with microtubules and the centrosome and is recruited to the IS of B cells: **A**. Scheme depicting activation of B cells with antigen-coated beads and representative confocal images of B cells labeled for actin (phalloidin) and Ecm29 activated for different time points (0, 30 and 60min). Dashed white circles represent the bead. **B**. Quantification of Ecm29 recruitment at the immune synapse (Bead) after different time points of activation. N=4. Cells>54. **C**. Scheme depicting activation of B cells on antigen-coated coverslips. Representative confocal images of centrin-GFP expressing B cells in resting (0 min) and activating (60 min) conditions, labeled for Ecm29 (magenta), microtubules, (α-tubulin, red). White arrows indicate the centrosome. Magnifications of centrosome areas (9μm^2^) are shown. **D and E**. Quantification of Ecm29 colocalization with microtubules and Ecm29 recruitment at the IS center in C. N=1. Cells>8. *p<0.05.****p<0.0001. One-way ANOVA with Tukey post-test and T-test was performed for all statistical analyses. Mean with SEM bars are shown. Scale bar = 10μm.

Next, we evaluated whether Ecm29 regulates the localization of the proteasome in B lymphocytes. For this purpose, we reduced the expression of Ecm29 in B cells by siRNA (**Figure EV2A and B**) and evaluated the distribution of the 26S proteasome in resting and activated B cells by immunofluorescence staining using an antibody that recognizes the 19S regulatory particle (RP). Upon Ecm29 silencing, we observed a reduction in the amount of proteasome at the centrosome of cells under resting conditions (**Figure 2A and C**). This result was confirmed by the detection of 19S RP by immunoblot in centrosome-rich fractions obtained from control and Ecm29-silenced cells. Furthermore, quantification of the 19S RP, normalized by γ-tubulin levels, indicated a reduction in the amount of the proteasome at the centrosome of Ecm29-silenced B cells (**Figure EV2E, F**). Moreover, quantification of proteasome activity in centrosome-rich fractions by using a fluorescent peptide as a substrate revealed a reduction in 50% of proteasome activity (**Figure EV2G**), which correlated with the decreased proteasome mass. When Ecm29-silenced cells were activated with antigen-coated beads, we observed enhanced accumulation of the proteasome at the antigen-bead contact site compare to control cells (blue area in line-scan) (**Figure 2A and B**). This result prompted us to further explore the distribution of the proteasome at the synaptic membrane of Ecm29-silenced B cells. To this end, we activated B cells by seeding them onto antigen-coated coverslips and labeled the 19S RP to visualize the proteasome as well as microtubules (α-tubulin), and the centrosome (Centrin-GFP). We found that the 19S RP significantly changed its distribution at the IS of Ecm29-silenced cells, exhibiting a more dispersed pattern, instead of colocalizing with microtubules (**Figure 2D and E**), suggesting that Ecm29 mediates the association of the proteasome with microtubules, analogously to what has been described in neurons (Hsu *et al*., 2015). Despite being more dispersed, the proteasome displayed slightly higher levels at the center of IS in Ecm29-silenced B cells (**Figure 2D and F**), suggesting that Ecm29 could be controlling proteasome distribution to the IS center, and potentially have an impact on the local degradation of protein targets within this region.

**Figure 2:**
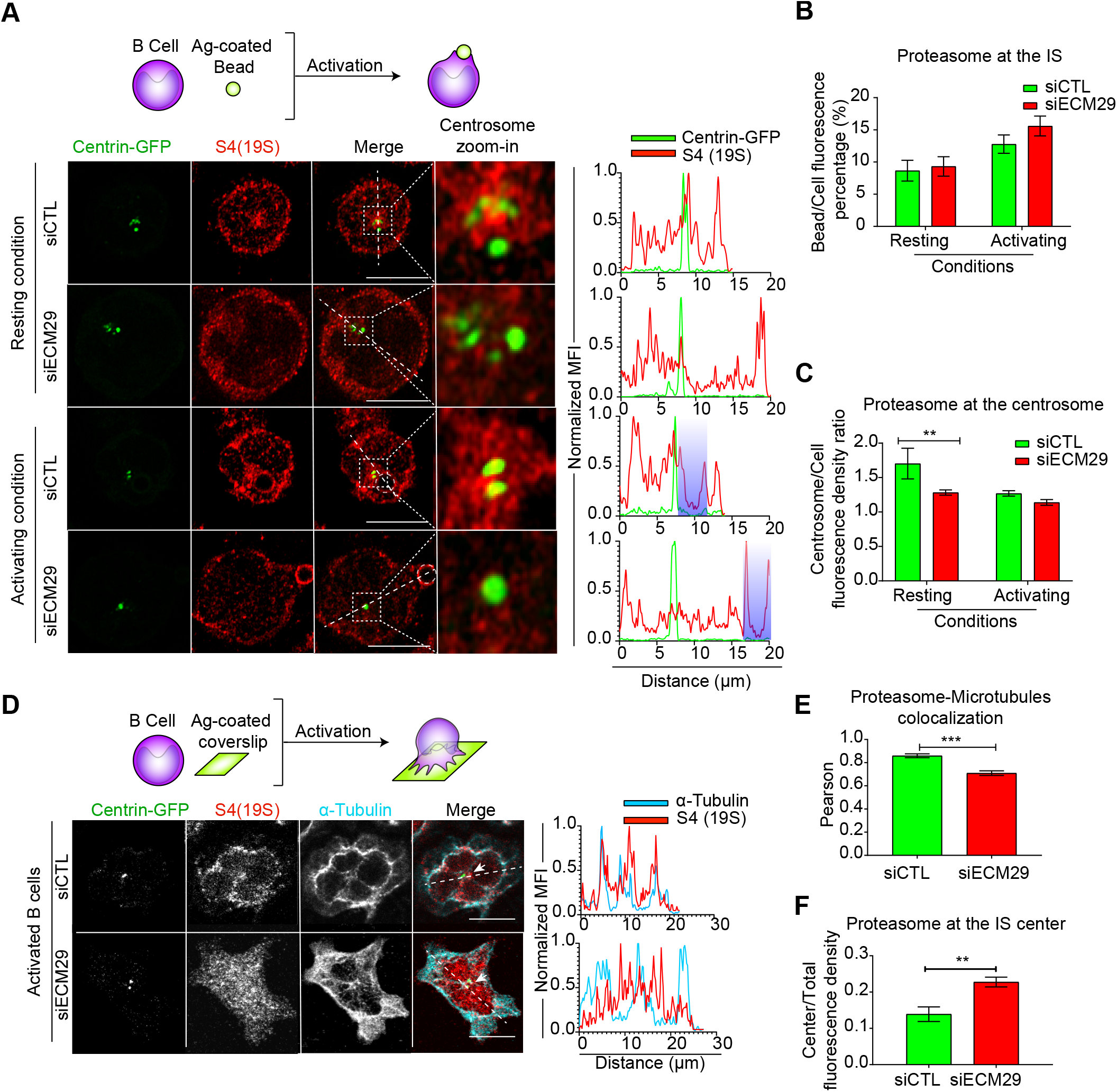
Ecm29 regulates proteasome localization at the centrosome and microtubules: **A**. Top: Schematic representation of B cells activated with antigen-coated beads. Representative confocal images of centrin-GFP expressing control (siCTL) and Ecm29-silenced (siECM29) B cells in resting (0 min) or activating (60 min) conditions, labeled for 19S[S4] (red). Magnifications of the centrosome area (3μm^2^) are shown. Fluorescence intensity distributions of centrin-GFP (green) and S4(19S) (red) across the cell (dashed white lines) are shown on the right. White dashed circles and shaded areas represent the bead and the centrosome area, respectively. **B and C**. Quantification of proteasome enrichment at the centrosome and the bead in A, respectively. N=1. Cells>41. **D**. Schematic representation of B cells activated on antigen-coated coverslips. Representative confocal images of centrin-GFP expressing control and Ecm29-silenced B cells after 60min of activation, labeled for 19S[S4] and α-tubulin. The fluorescence intensity distributions of α-tubulin (blue) and S4(19S) (red) across the cell (dashed white lines) are shown. White arrows indicate the centrosome. **E and F**. Quantification of proteasome colocalization with microtubules and proteasome recruitment at the IS center, respectively. N=1.Cells>7. **p<0.01. ***p<0.001. Two-ways ANOVA with Sidak’s post-test and T-test was performed for all statistical analyses. Mean with SEM bars are shown. Scale bar = 10μm.

To characterize the dynamic recruitment of the proteasome at the IS in control versus Ecm29-silenced cells, we labeled the proteasome in live cells by using a specific fluorescent probe that binds to the catalytic β5 subunit (Bsc2118-FL-Bodipy). We used this probe at 5nM in our assays, previously described not to significantly inhibit proteasome activity (Mlynarczuk-Bialy *et al*., 2014). Additionally, we verified that this dose did not generate an accumulation of ubiquitylated proteins (**Figure EV3A**), suggesting no significant effects over proteasome activity. Using this approach, we confirmed that the proteasome co-distributed with microtubules, displaying a central and peripheral localization at the synaptic membrane (**Figure EV3B**), with limited diffusion rates (median: 0.074 μm^2^/sec) (**Figure EV3C and D**), comparable to those described in neurons (Otero *et al*., 2014). Using this experimental approach, we also verified that in Ecm29-silenced cells, the 20S proteasome became more enriched at the IS than control counterparts, which was evidenced by a higher number and duration of proteasome tracks (**Figure EV4A, C, and E**). When measuring proteasome diffusion rates, we observed that these were slightly higher in Ecm29-silenced cells without affecting their overall displacement at the IS (**Figure EV4A, B, and D**). Altogether, these results suggest that Ecm29 regulates the recruitment of the 20S proteasome to the IS, controlling its diffusion rate, most likely in a microtubule-dependent manner.

Ecm29 has been shown to negatively regulate the proteasome by inhibiting its ATPase activity, which is crucial to unfold and translocate target proteins into the catalytic core for degradation (De La Mota-Peynado *et al*., 2013). To determine whether Ecm29 silencing affected the activity of the proteasome, we measured the amount of ubiquitylated proteins and proteasome subunits in Ecm29-silenced and control B cells. Quantification by immunoblot of proteasome subunits: S4 for the regulatory particle (19S), α-β for the catalytic core (20S), and LMP7 for the induced catalytic core (immune proteasome), showed no differences between control and Ecm29-silenced cells (**Figure EV2C and D**). These results indicate that in B cells, Ecm29 regulates the localization of the proteasome without significantly affecting its total mass or activity.

### Ecm29 regulates actin remodeling at the synaptic membrane and B cell spreading responses

Having shown that Ecm29 regulates the distribution of the proteasome in B cells and considering that proteasome activity is crucial for actin remodeling at the centrosome and IS of B cells (Ibañez-Vega, Del Valle Batalla, *et al*., 2019), we sought to determine whether actin levels at these two regions were affected by Ecm29 depletion. To this end, we seeded control and Ecm29-silenced B cells onto antigen-coated coverslips and imaged them by confocal microscopy, measuring the spreading area, delimited by actin labeling. Unexpectedly and contrary to the effect of inhibiting proteasome activity (Ibañez-Vega, Del Valle Batalla, *et al*., 2019), we found that, after 30 and 60 min of activation, Ecm29-silenced cells displayed an increased spreading area, which was at least two-fold higher than control cells (**Figure 3A and B**). We next characterized actin dynamics at the synaptic interface of these cells by performing live imaging using TIRFM. For this purpose, control and Ecm29-silenced B cells expressing the actin fluorescent reporter LifeAct-mCherry were activated onto antigen-coated coverslips. Similar to our observations in fixed cells, Ecm29-silenced cells displayed an increased spreading response during activation, which was also approximately two-fold higher than control cells (**Figure 3C and D**). Under these conditions, we noticed that the spreading rate was also higher in Ecm29-silenced B cells (**Figure 3D**). Interestingly, these cells underwent continuous spreading, which was not followed by a stationary phase, generally observed in control cells after 10 min of activation, where spreading starts to slow down (**Figure 3D**). This observation suggests that Ecm29-silenced B cells sustain an uncontrolled spreading response without reaching a stationary phase or further contraction, as was previously shown by B cells interacting with membrane-tethered antigens (Fleire *et al*., 2006).

**Figure 3:**
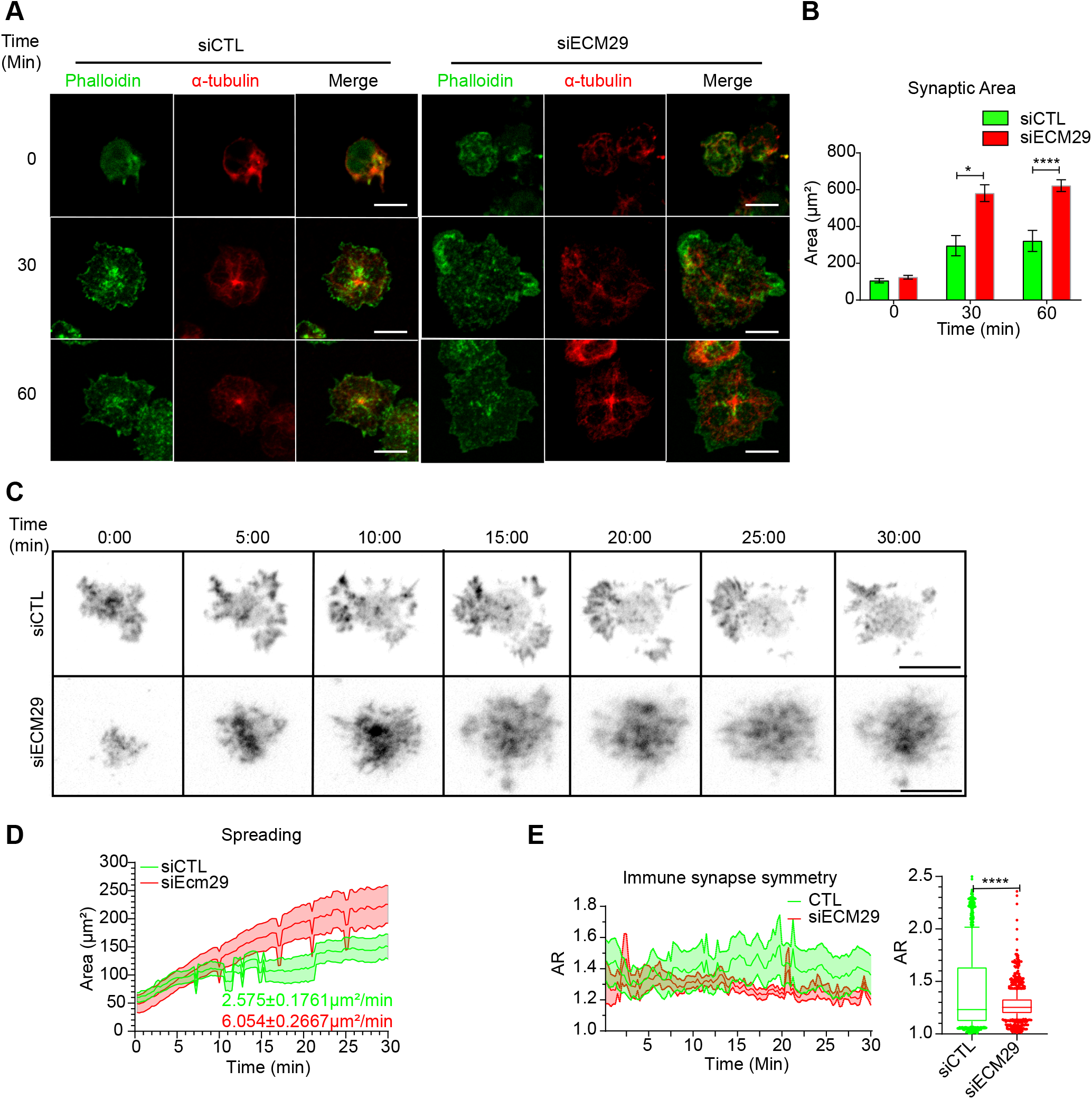
Ecm29 silencing enhances B cell spreading responses: **A**. Representative confocal images of control (siCTL) and Ecm29-silenced (siECM29) B cells activated for different time points (0, 30, and 60 min), stained with phalloidin (green) and α-Tubulin (red) are shown. **B**. Quantification of the spreading area in A. N=3. Cells>40. **C**. Representative Time-lapse images acquired byTIRFM of control and Ecm29-silenced B cells expressing LifeAct-mCherry (greyscale). **D and E**. Quantification of the spreading area and aspect ratio (AR) in C, respectively. The mean spreading velocity and mean AR are shown. N=10. *p<0.05.****p<0.0001. Two-way ANOVA with Sidak’s post-test and T-test were performed. Mean with SEM bars (B), lines (D and E), and boxes and whiskers with 10% percentile (E) are shown. Scale bar=10μm.

In T lymphocytes, immune synapse stability has been associated with forming symmetric synapse (Kumari *et al*., 2020). Given that Ecm29-silenced B cells displayed an uncontrolled spreading response, we investigated whether this was due to perturbed immune synapse stability. To this end, we quantified the IS symmetry during the spreading response in control and Ecm29-silenced B cells, measured as the aspect ratio of the spreading area. We found that the symmetry of the synapse in Ecm29-silenced cells was highly sustained compared to control cells (**Figure 3E**), suggesting that Ecm29 silencing enhances immune synapse stability in B cells.

Ecm29-silenced B cells also exhibited smaller lamellipodia, which could result from defects in actin polymerization, required to generate actin retrograde flow (Wang and Hammer, 2020). Thus, we sought to determine whether actin polymerization at lamellipodia, measured as the velocity of actin-retrograde flow, was also affected. For this purpose, we seeded LifeAct-mCherry expressing control and Ecm29-silenced B cells for 30 min in antigen-coated coverslips before performing live-cell imaging. We observed that the length of lamellipodia was significantly reduced in Ecm29-silenced B cells, which also showed a reduction in the velocity of actin-retrograde flow, compared to control cells (**Figure 4A and B**). Altogether these results highlight a role for Ecm29 in forming lamellipodia at the IS of B cells, and suggests that the proteasome close to the IS membrane could be regulating actin polymerization, probably, by the degradation of actin polymerizing factors, as was previously suggested (Hao *et al*., 2013; Ibañez-Vega, Del Valle Batalla, *et al*., 2019).

**Figure 4:**
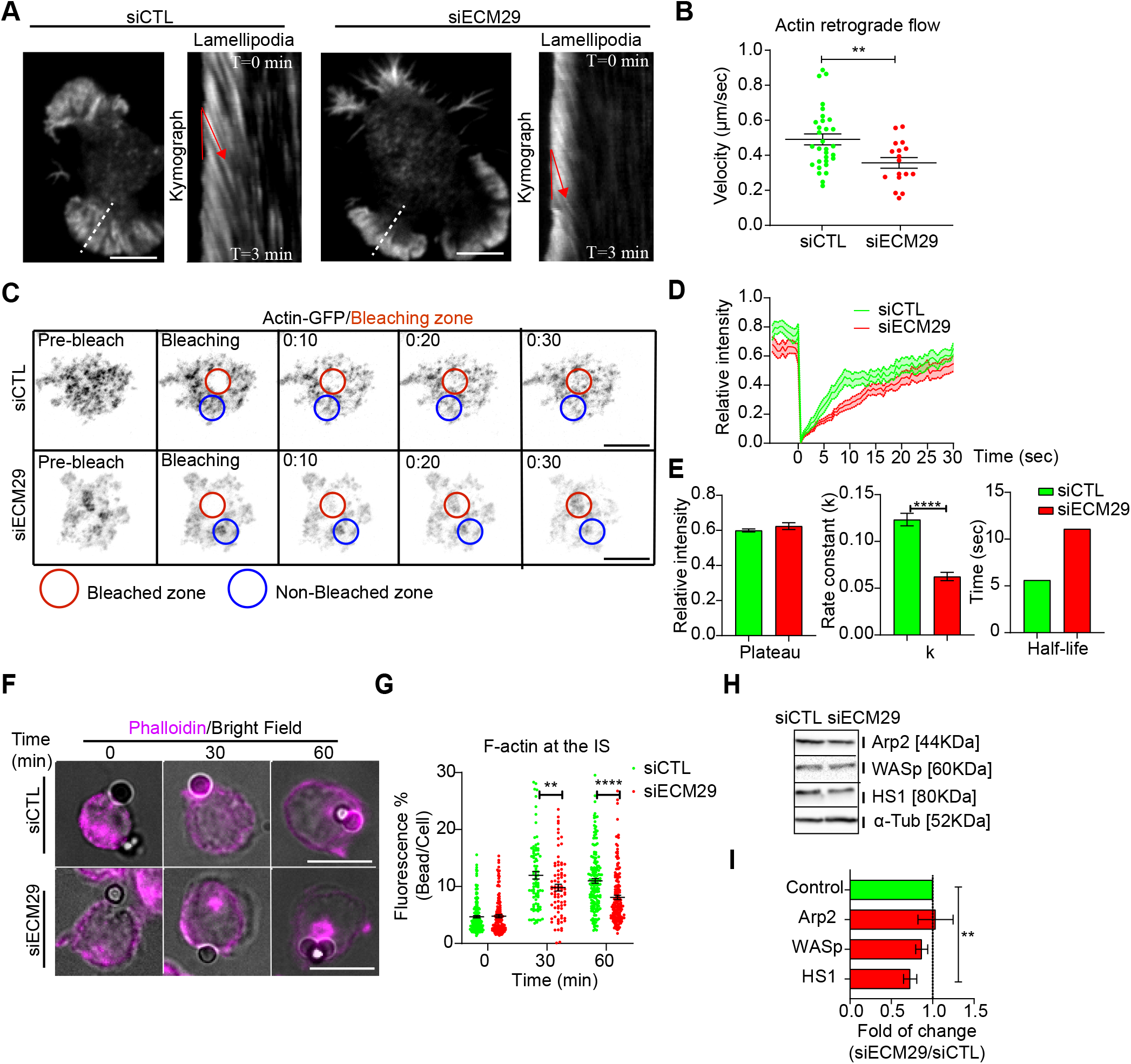
Ecm29 silencing reduces actin dynamic and accumulation at the IS: **A**. Representative TIRFM images of control (siCTL) and Ecm29-silenced (siECM29) B cells after 30 min of activation and their respective kymographs of lamellipodia (white dashed line). Red lines represent the actin retrograde flow angle. **B**. Quantification of actin retrograde flow in A. N=17. **C.** Time-lapse FRAP-TIRFM images of control and Ecm29-silenced B cells expressing Actin-GFP (greyscale) after 30 min of activation. Red and blue circles represent the bleached and non-bleached zone, respectively. **D and E**. Quantification of Fluorescence Recovery After Photobleaching of Actin-GFP, and associated parameters (plateau, k, and half-life). N=15. **F**. Representative epifluorescence images of control and Ecm29-silenced B cells activated with antigen-coated beads for different time points, labeled for F-actin (Phalloidin) are shown. **G.** Quantification of F-actin accumulation at the bead in F. N>3. Cells>80. **H.** Representative immunoblot of control and Ecm29-silenced B cells. Arp2, WASp, HS1, and α-Tub are shown. **I**. Quantification of protein levels in E. N>4. **p<0.01). ****p<0.0001. A T-test was performed. Mean with SEMs and individual experiments (points) (A and G), and boxes with SEM (I) are shown. Scale bar=10μm.

To further evaluate the impact of Ecm29 on actin polymerization at the IS, we seeded actin-GFP-expressing B cells, silenced or not for Ecm29, onto antigen-coated coverslips for 30 min and performed a Fluorescence Recovery After Photobleaching (FRAP) assay using TIRFM. This assay allowed us to quantify actin turnover at the synaptic interface, as an indirect measurement of actin polymerization, previously reported in B cells (Pauls, Hou and Marshall, 2020). Indeed, Ecm29-silenced cells showed a reduction in fluorescence recovery velocity, expressed as the rate constant (k) (**Figure 4C, D, and E**), suggesting that actin polymerization at the synapse is reduced under these conditions. To determine whether reduced actin polymerization at the IS in Ecm29-silenced cells translated into less F-actin accumulation at the synaptic interface, we activated B cells with antigen-coated beads, measuring its accumulation at the antigen contact site. As expected, after different time points of activation, we observed reduced F-actin accumulation at the antigen contact site (bead) in Ecm29-silenced cells compared to control cells (**Figure 4F and G**). Considering that Ecm29-silenced cells accumulate more proteasome at the IS, this highlights its negative correlation with F-actin accumulation at the IS.

To further explore a functional link between proteasome and actin polymerization at the IS, we co-labeled F-actin (LifeAct-mCherry) and the 20S proteasome (5nM Bsc2118-FL-Bodipy) in live cells activated on antigen-coated coverslips for 30 min and analyzed the synaptic interface by TIRFM. We found that structures labeled for the proteasome negatively correlated with Factin fluorescence (**Figure EV5A-F, and I**). This negative correlation was dependent on proteasome activity because pre-treatment with 5μM MG-132 or high concentrations of Bsc2118-FL-Bodipy (100nM), which inhibits proteasome activity, abolished this negative correlation. On the other hand, the silencing of Ecm29, did not affect this negative correlation (**Figure EV5G, H and I**). Altogether, these results suggest that the proteasome at the synaptic interface is associated with F-actin depletion, with functional repercussions in actin turnover and lamellipodia formation.

To determine how the proteasome negatively regulates actin accumulation at the IS of B cells, we evaluated whether the levels of actin polymerizing factors, which are targets of proteasome degradation (Schaefer, Nethe and Hordijk, 2012), changed in Ecm29-silenced cells. The levels of actin polymerizing factors, Arp2, WASp, and HS1, which have been shown to play a role in B cells, (Bolger-Munro *et al*., 2019; Roper *et al*., 2019) were quantified in control and Ecm29-silenced B cells, by immunoblot. We found reduced levels of HS1 in Ecm29-silenced cells, whereas Arp2 or WASp remained unchanged (**Figure 4H and I**). Considering that HS1 is localized at the synaptic interface and the centrosome in B cells (Obino *et al*., 2016), our results suggest that, HS1 could be the main target of proteasome degradation at the synaptic interface during B cell activation.

### Ecm29 silencing enhances BCR clustering and signaling

Having shown that Ecm29 regulates actin polymerization at the IS, we next explored the role of Ecm29 in BCR signaling. F-actin depolymerization facilitates BCR diffusion at the IS, promoting receptor clustering, and signaling (Freeman et al., 2015; Tolar, 2017). Given that Ecm29 silencing reduced actin accumulation and turnover at the synapse, we evaluated whether the distribution of the BCR was affected under these conditions. Our results show that Ecm29-silenced cells displayed enhanced BCR accumulation at the center of the IS, especially after 30 min of activation, shown by the higher mean fluorescence (MFI) this region (the first quartile) (**Figure 5A and B**). This also translated into higher BCR downstream signaling where Erk phosphorylation levels, were higher in Ecm29-silenced cells compared to controls (**Figure 5C**). Thus, enhanced BCR clustering and signaling could result from unstable actin structures at the IS, increasing BCR mobility, as previously described (ref PMID: 20171124, Tolar, 2017). These observations suggest that the reduced actin polymerization at the IS in Ecm29-silenced B cells results in enhanced B cell activation. Considering that actin cytoskeleton remodeling is also important at the centrosome for antigen extraction and presentation (Obino *et al*., 2016; Ibañez-Vega, Del Valle Batalla, *et al*., 2019), we next evaluated whether Ecm29 silencing could affect centrosome repositioning upon BCR activation.

**Figure 5:**
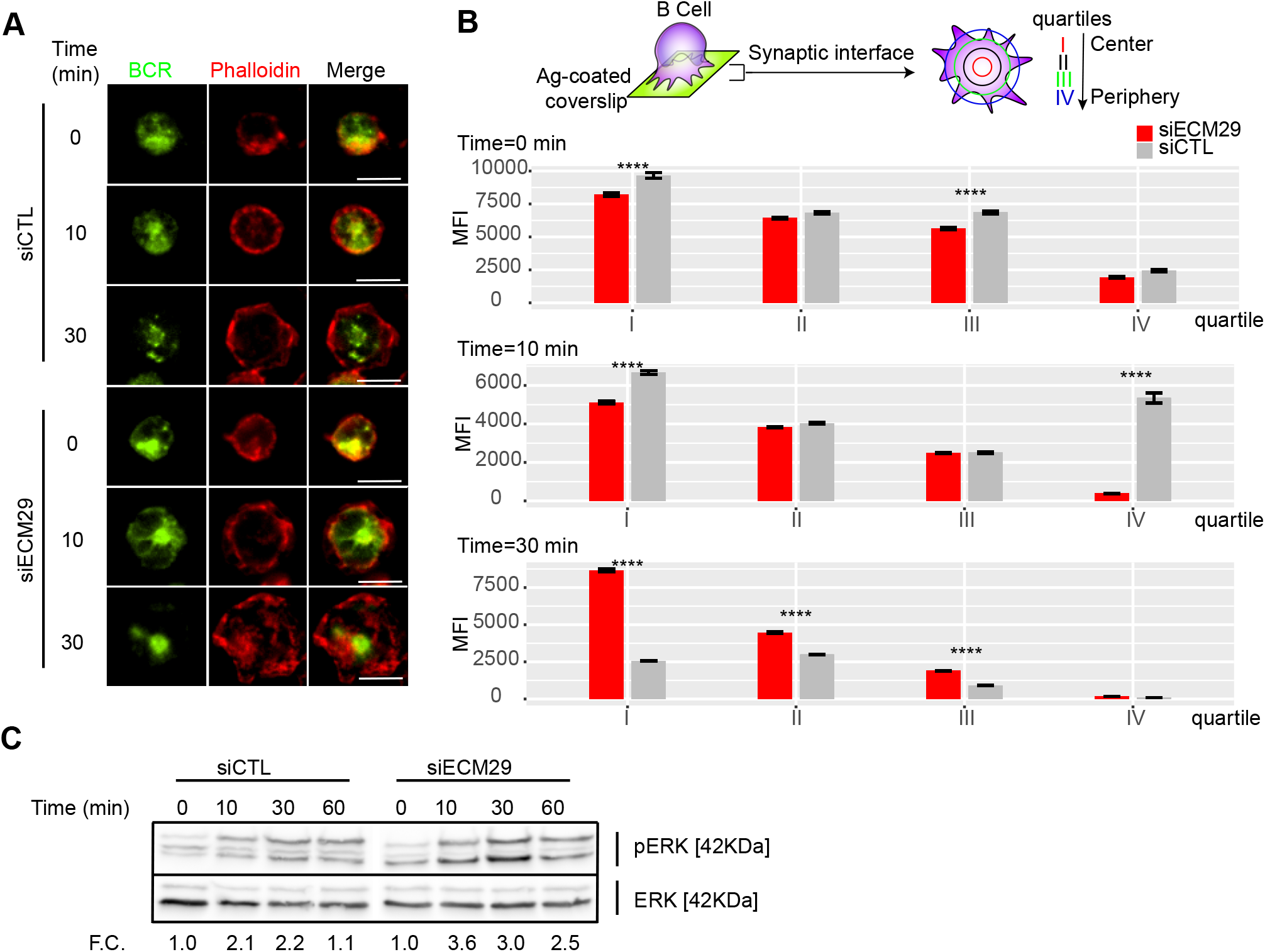
Ecm29 silencing enhances BCR clustering and Erk phosphorylation: **A**. Representative Epifluorescence images of control (siCTL) and Ecm29-silenced (siECM29) B cells, activated onto antigen-coated coverslips for different time points (0, 10 and 30 min), labeled for the BCR and F-actin (phalloidin). **B**. Schematic representation of BCR distribution quantification, and the respective quantification of BCR accumulation in each quartile of the immune synapse in B cells shown in A. N=2. Cells>30. **C**. Immunoblot of protein of control and Ecm29-silenced B cells activated for different time points (0,10, 30, and 60 min). Phosphorylated Erk (pErk) and total Erk (Erk) are shown. F.C.= Fold of change respect time 0 min. *p<0.05. two-way ANOVA with Sidak’s post-test. Scale Bar= 10μm.

### Ecm29 regulates centrosome positioning at the IS

A hallmark of the B cell IS is the repositioning of the centrosome to the synaptic interface, which orchestrates the recruitment of lysosomes to the IS, where they undergo local secretion to facilitate antigen extraction (Yuseff *et al*., 2011). Centrosome repositioning to the synaptic interface requires depolymerization of perinuclear actin that maintains the centrosome linked to the nucleus (Obino *et al*., 2016). We previously described that proteasome activity is crucial for actin depletion at the centrosome to enable centrosome repositioning (Ibañez-Vega, Del Valle Batalla, *et al*., 2019). Thus, we asked whether silencing Ecm29 in B cells had a similar effect, given that it results in the mislocalization of the proteasome. For this purpose, we activated control and Ecm29-silenced B cells expressing Centrin-GFP on antigen-coated coverslips and measured the recruitment of the centrosome to the synapse after different time points of activation. As previously described, we found that the centrosome reached the proximity of the antigen-coated surface in control conditions, which we detected at the first and second Z-axis section of the fluorescence distribution graph (**Figure 6A and B**). Conversely, in Ecm29-silenced cells, the centrosome was not recruited to the IS, where the mean fluorescence of Centrin-GFP remained between the third and sixth Z fraction of the fluorescence distribution graph (**Figure 6A and C**). Next, we quantified the amount of actin at the centrosome in control and Ecm29-silenced cells. Our imaging analysis showed that Ecm29-silenced B cells exhibit defective actin clearance at the centrosome (**Figure 6D and E**), which most likely results from a decrease in proteasome activity associated with the centrosome of these cells (**Figure 2A, B, and EV2**).

**Figure 6:**
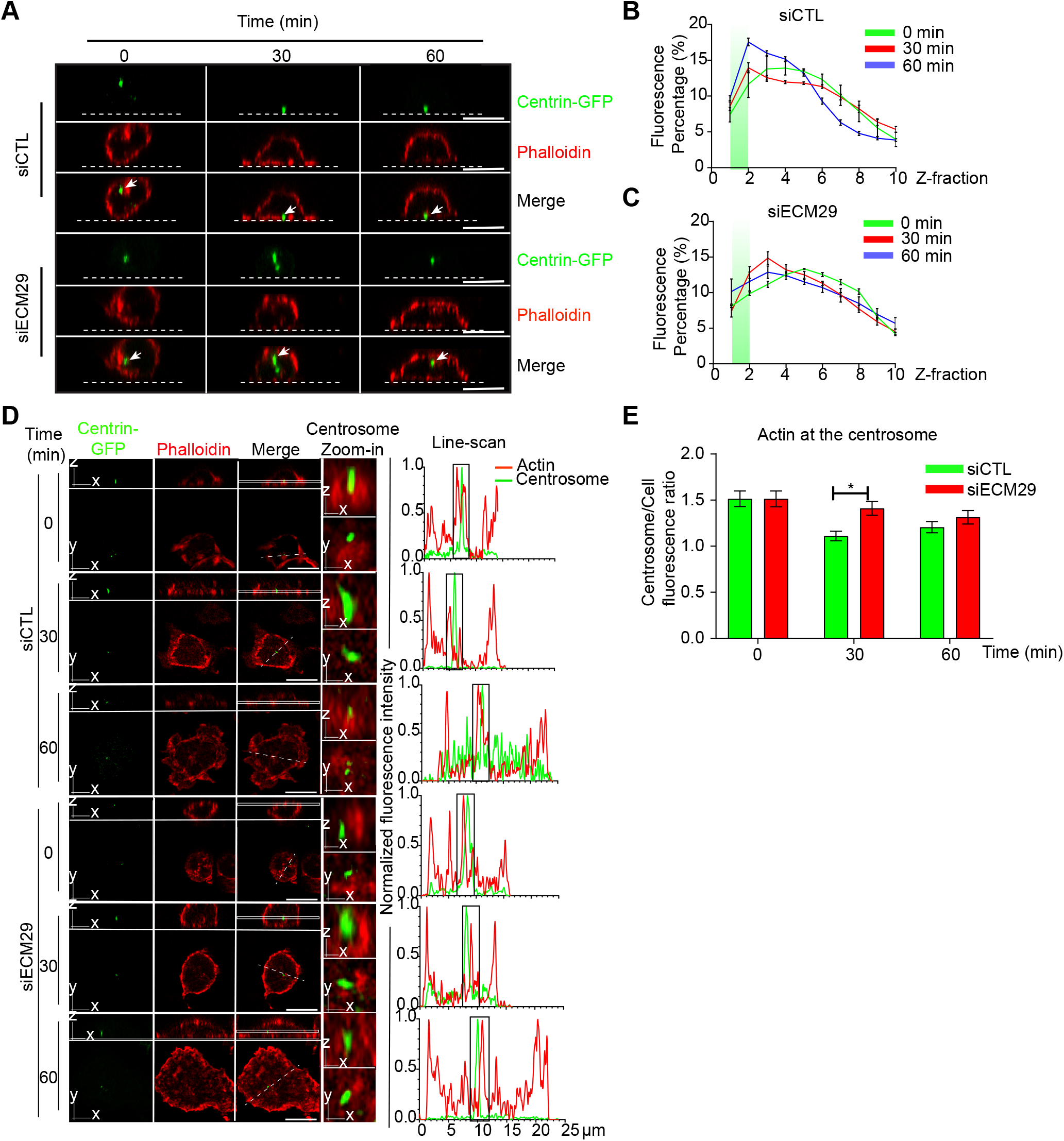
Centrosome polarization and perinuclear actin clearance rely on Ecm29: **A**. Representative X/Z confocal images of centrin-GFP expressing control (siCTL) and Ecm29-silenced (siECM29) B cells activated on antigen-coated coverslips for different time points, stained for F-actin, phalloidin are shown. White arrows indicate the centrosome. The white dashed line represents the position of the coverslip. **B and C**. Quantification of centrosome fluorescence intensity along the Z dimension from the coverslip to the upper cell limit of control and Ecm29 silenced B cells, respectively. The green rectangle represents the synaptic Z area (between 1 and 2 Z-fraction), and the maximum value of the curve represents the localization of the centrosome in the Z dimension. N=1. Cells>3. **D**. Representative Ayriscan images of centrin-GFP expressing control and Ecm29-silenced B cells activated on antigen-coated coverslips for different time points, labeled for F-actin, phalloidin. Magnifications of the centrosome (9μm^2^) and the fluorescence intensity distribution of centrin-GFP (green) and phalloidin (red) across the cell (dashed white lines) are shown. White rectangles represent the X/Y position for images in Z. **E**. Quantification of actin at the centrosome in D. N=2. Cells>45. *p<0.05. Two-ways ANOVA with Sidak’s post-test and T-test was performed for all statistical analyses. Mean with SEM lines (B and C) and bars (E) are shown. Scale bar = 10μm.

To confirm our results, we measured the distance between the centrosome and the nucleus in control and Ecm29 silenced B cells activated with antigen-coated beads. As expected, in Ecm29-silenced cells the centrosome remained opposed to the nucleus, confirming their deficiency to translocate their centrosome to the synaptic membrane (**Figure EV6A, B, and C**).

### Ecm29 silencing impairs antigen extraction and presentation in B cells

Defective centrosome repositioning to the IS impairs the recruitment and local secretion of lysosomes at the synaptic membrane, which can facilitate the extraction and processing of immobilized antigens (Yuseff *et al*., 2011; Ibañez-Vega, Del Valle Batalla, *et al*., 2019). Therefore, we investigated the functional consequences of Ecm29 silencing on antigen extraction and presentation by B cells. To this end, we performed an antigen presentation assay using control or Ecm29-silenced B cells (**Figure 7A and B**). Our results show that B cells silenced for Ecm29 display defective antigen presentation and reduced cell surface expression of MHC-II, as evidenced by lower levels of loaded peptide presentation by Ecm29-silenced cells to T cells (**Figure 7B**). The defects in antigen presentation could result from impaired antigen extraction and possibly MHC-II trafficking to the cell membrane. To evaluate this possibility, we activated control or Ecm29 silenced B cells with OVA-antigen-coated beads for different time points and measured the amount of OVA fluorescence remaining on the beads as an indicator of antigen extraction. We found that Ecm29 silenced cells showed higher OVA-antigen levels on beads after activation compared to control cells, confirming that Ecm29-silenced B cells could not efficiently extract antigen (**Figure 7C and D**). Considering that antigen extraction relies on lysosome recruitment and secretion (Yuseff *et al*., 2011), we also followed the distribution of lysosomes labeled for Lamp1 in activated B cells and noticed that Ecm29 displayed delayed recruitment of lysosomes to the IS (**Figure 7D and E**). Overall, these results suggest that lysosome trafficking to the IS depends on Ecm29, which promotes centrosome repositioning by depolymerizing actin within this region.

**Figure 7:**
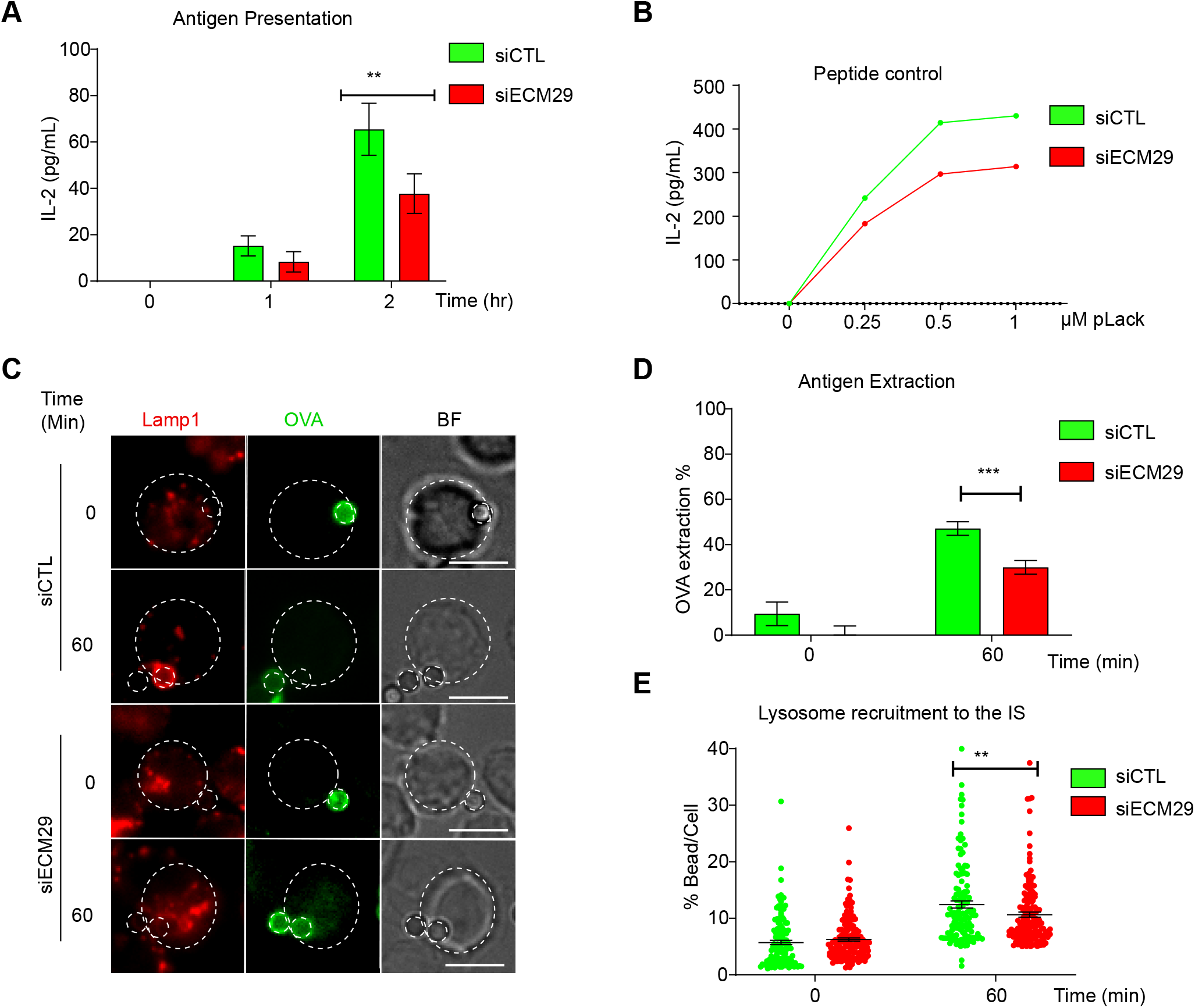
Efficient antigen extraction and presentation requires Ecm29: **A**. Antigen presentation assay for control (siCTL) and Ecm29 silenced (siECM9) B cells. Levels of IL-2 by T cells were quantified by ELISA. N=3. **B**. Representative graph of peptide controls for cells used in antigen presentation assays. **C**. Representative images of control and Ecm29 silenced B cells activated with beads coated with anti-IgG+OVA in resting (0 min) and activating conditions (60 min). Lysosomes (Lamp1) and antigen (OVA) are shown. The white dashed circles delimit cell boundaries and bead. **D and E**. Quantification of antigen extraction, measured as the amount of OVA extracted from the bead, and lysosome recruitment to the bead in C. N=4. Cells>82. **p<0.01. ***p<0.001. Two-ways ANOVA with Sidak’s post-test and T-test was performed for all statistical analyses. Mean with SEM bars (A and D), individual cells (points) (E) are shown. Scale bar = 10μm.

In summary, we put forward a novel mechanism involved in regulating the B cell immune synapse, where Ecm-29 dependent proteasome distribution orchestrates actin remodeling at the synaptic interface and the centrosome, thereby controlling key cellular functions such as lysosome trafficking and antigen extraction and presentation.

## Discussion

A functional immune synapse relies on actin remodeling at the synaptic membrane and the centrosome: The former promotes BCR clustering and downstream signaling (Freeman et al., 2015; Spillane & Tolar, 2018; Tolar, 2017), and the latter enables centrosome repositioning to the IS together with lysosomes, which, upon secretion, facilitate antigen extraction (Yuseff *et al*., 2011; Obino *et al*., 2016; Ibañez-Vega, Del Valle Batalla, *et al*., 2019). Our work reveals that the localization of the proteasome at the synaptic interface and centrosome relies on Ecm29, which in turn regulates actin remodeling in both compartments; and, therefore, plays a pivotal role in immune synapse formation, antigen extraction, and presentation.

A role of Ecm29 in regulating the localization of the proteasome has been previously reported in neurons (Otero *et al*., 2014; Lee *et al*., 2020), but has remained relatively unexplored in lymphocytes. The underlying mechanisms by which Ecm29 regulates the intracellular localization of the proteasome include: 1) promoting the interaction of the proteasome with motor proteins, such as kinesins and dyneins, which directly anchors the proteasome to microtubules, and is responsible for proteasome retrograde and anterograde movement (Hsu *et al*., 2015) 2) mediating the association of the proteasome to vesicles and favors its fast movement by hitch-hiking (Otero *et al*., 2014) and 3) promoting the association of the proteasome with Rab11+ recycling vesicles and organelles, such as Endoplasmic Reticulum and the centrosome (Gorbea *et al*., 2004, 2010). Our work suggests that Ecm29 mediates the association of the proteasome to microtubules and the centrosome in B cells; however, the molecular mechanisms involved in mobilizing the proteasome remain to be explored.

Interestingly, Ecm29 has also been described to act as a proteasome inhibitor and quality control protein, where its association to the proteasome inhibits the ATPase activity of the regulatory particle and stabilizes its interaction with the catalytic core (Lehmann *et al*., 2010; De La Mota-Peynado *et al*., 2013; Haratake *et al*., 2016). However, we found that Ecm29 silencing did not affect overall levels of proteasome activity, measured as the accumulation of ubiquitylated proteins, which suggest that in B cells, Ecm29 regulates proteasome localization rather its activity. Despite of this, further analysis of how Ecm29 affects specific proteasome functions, such as degradation rate, ATPase activity, or specific protease activity, are needed to completely discard whether or not Ecm29 affects proteasome activity.

B cells silenced for Ecm29 displayed lower levels of proteasome at the centrosome, which correlated to increased F-actin at this level, suggesting that centrosome-associated proteasome could act as a negative regulator of actin polymerization within this region. This observation is consistent with our previous findings, where inhibition of proteasome activity also induced an accumulation of F-actin at the centrosome and defective actin clearance upon B cell activation, disabling centrosome repositioning to the synapse (Obino *et al*., 2016; Ibañez-Vega, Del Valle Batalla, *et al*., 2019). In the same line, our results show that decreasing the proteasome at the centrosome, by silencing Ecm29, also leads to defective centrosome repositioning at the IS. As anticipated, defective centrosome polarization in Ecm29 silenced B cells correlated with poor lysosome recruitment to the IS and impaired antigen extraction and presentation. Thus, our results shows that the proteasome pool associated to the centrosome is crucial for centrosome proteostasis, which is concordance with previous observations that suggest that the centrosome as a platform of local UPS-mediated degradation (Vora and Phillips, 2016).

A striking observation reported here is that upon activation, Ecm29-silenced cells accumulated higher proteasome levels at the cortical and synaptic regions. Thus, it is tempting to speculate that Ecm29 favors the centrosomal localization of the proteasome, excluding it from the cell cortex and the IS. The molecular mechanisms underlying the cortical localization of the proteasome are not understood in lymphocytes. However, in neurons, adaptor proteins, such as GPM6A/B, were shown to mediate the interaction of the proteasome with the plasma membrane (Ramachandran and Margolis, 2017). Thus, it would be interesting to address whether the mechanisms involved in the localization of cortical proteasome in neurons is shared by lymphocytes, and how the proteasome controls its association with proteasome-regulators.

In addition to enhanced proteasome levels at the IS, Ecm29 silenced cells displayed reduced actin turnover and slower actin-retrograde flow at the synaptic interface, suggesting that actin polymerization at the IS was reduced within this zone. These observations are consistent with the general view of the proteasome as a negative regulator of actin polymerization (Schaefer, Nethe and Hordijk, 2012; Hsu *et al*., 2015), where actin nucleating factors are selectively downregulated by the ubiquitin-proteasome system (de la Fuente *et al*., 2007), thereby turning down actin polymerization. This idea is further supported by our observations, where we found that HS1 was decreased upon Ecm29 silencing, indicating that this protein could be targeted for proteasome degradation. Indeed, HS1 has five ubiquitin sites (K34, K60, K123, K192, and K239) (Hornbeck *et al*., 2012). In B cells, BCR activation triggers HS1 phosphorylation by Syk, and its subsequent recruitment to the IS, being crucial in the promotion of actin assembly at the IS (Hao, Carey and Zhan, 2004; Obino *et al*., 2016). Thus, it is possible that silencing Ecm29, which results in increased levels of proteasome at the IS and enhanced HS1 degradation, could be responsible for poor actin accumulation at the synaptic interface, upon BCR activation. On the other hand, Ecm29 silencing did not alter WASp levels, which could be a consequence of its interaction with the WASp interacting protein (WIP), which protects WASp from degradation (de la Fuente *et al*., 2007). Indeed, WIP is also recruited to the IS of B cells (Keppler *et al*., 2018), supporting the idea that WIP could be protecting WASp from degradation in Ecm29-silenced B cells.

The actin cytoskeleton plays a critical role in BCR signaling by restricting its lateral diffusion and association with co-receptors (Tolar, 2017). Indeed, B cells treated with drugs that disrupt actin organization induce BCR signaling in the absence of ligand (Batista, Treanor and Harwood, 2010). Additionally, proteins belonging to the Ezrin-Radixin-Moesin Family (ERM family), which link the actin cytoskeleton to the plasma membrane, negatively regulate BCR lateral diffusion (Treanor *et al*., 2009; Liu *et al*., 2013; Tolar, 2017a). Interestingly, ezrin also can be degraded by the proteasome (Grune *et al*., 2002; Suetsugu *et al*., 2002; Sun *et al*., 2019) and it has been reported that upon BCR activation, an early ubiquitylation response affects the BCR downstream kinases, signaling components, such as LAT2, RAC1, CDC42, VAV1 and Ezrin (Satpathy *et al*., 2015). Thus, in addition to actin polymerizing factors, the proteasome at the IS could be degrading proteins involved in BCR activation, as well as other factors that couple actin to the cell cortex, such as ezrin, and control membrane tension (Le Bouteiller *et al*., 2011; Schaefer, Nethe and Hordijk, 2012; Ingber, Wang and Stamenović, 2014; Kelkar, Bohec and Charras, 2020). Consequently, the larger spreading area displayed by Ecm29 silenced B cells, which accumulate more proteasome at the synaptic membrane, could result from a relaxed cortex or enhanced BCR signaling. Thus, the timely recruitment of the proteasome to the immune synapse is crucial to determine where and which proteins would be degraded.

Interestingly, microtubules drive the spreading response in fibroblasts, where the rapid growth of microtubules toward the cell borders is essential for isotropic spreading (Tvorogova *et al*., 2018). In our study, we found that Ecm29-silenced B cells displayed a highly symmetric IS, which resembles an isotropic spreading response. Thus, it is tempting to speculate that the increased spreading response and the slow actin-retrograde flow in Ecm29-silenced B could allow microtubule growth toward the cell margins, which would lead to a sustained spreading and reduced actin-retrograde flow. Indeed, an analogous negative correlation between the actin cytoskeleton and microtubules was previously described at the centrosome, where the reduction of polymerized actin at the centrosome triggered an increased microtubule growth (Inoue1 *et al*., 2019). The role of the proteasome selectively degrading molecules that tune the microtubule-actin crosstalk at the synaptic membrane shall provide insights on how antigen extraction and processing are regulated at the B cell synapse.

In conclusion, our work reveals that the distribution of the proteasome, mediated by Ecm29, controls the formation of the IS by regulating actin dynamics at the centrosome and synaptic membrane. These new findings contribute to understanding how B lymphocytes efficiently manage to orchestrate complex actin cytoskeleton remodeling at these two levels and control the establishment of a polarized phenotype during IS formation.

## Materials and methods

### Cell lines and culture

The mouse lymphoma cell line IIA1.6, which is a FcγR-defective B cell line with the phenotype of quiescent mature B-cells (Lankar *et al*., 2002) and the LMR7.5 Lack T-cell hybridoma, which recognizes I-Ad-LACK_156–173_ complexes, were cultured as previously described (Vascotto *et al*., 2007) in CLICK medium (RPMI 1640, 10% fetal bovine serum, 100U/mL penicillin-streptomycin, 0.1% β-mercaptoethanol, and 2% sodium pyruvate. For proteasome inhibition, 5×10^6^ B cells/mL were incubated with 5μM MG-132 for 1 hour at 37°C before functional analysis.

### Antibodies and reagents

We used rat anti-mouse LAMP1 (BD Bioscience, #553792, 1:200), rabbit anti-mouse α-Tubulin (Abcam, ab#6160, 1:200), rabbit anti-acetyl-α-Tubulin (Lys40; cell signaling, #5335, 1:200), rabbit anti-mouse γ-Tubulin (Abcam, #Ab11317, 1:1000), rabbit antimouse S4/19S RP (Abcam, #Ab223765, 1:100), rabbit anti-mouse αβ/20S proteasome (Abcam, #Ab22673, 1:200), anti-mouse Ecm29 (Abcam, #Ab28666, 1:100), mouse anti-mouse Ubiquitin P4D1 (Santa Cruz, #Sc-8017, 1:1000), anti-mouse anti-actin (cloneC4, ImmunO, #691001), rabbit anti-mouse Arp2 (Cellsignal, #5614, 1:500), goat anti-mouse IgGFab^2^ (Jackson ImmunoResearch), rabbit anti-OVA (Sigma-Aldrich, #C6534, 1:500). For secondary antibodies: donkey anti-rabbit IgG-Alexa488 (LifeTech, 1:200), goat anti-rabbit IgG-Alexa546 (ThermoScientific, 1:200), Donkey anti-rat IgG-Alexa546 (ThermoScientific, 1:200), Donkey anti-rat-Alexa647 (ThermoScientific, 1:200), Phalloidin-Alexa-647 (Life Technology, #22287, 1:200), DAPI (Abcam). Ovalbumin was purchased from Sigma-Aldrich, MG-132, and Epoxomicin were purchased from Merk (Millipore). Bsc2118-FL-Bodipy was kindly provided by Ulrike Kuckelkorn (Mlynarczuk-Bialy *et al*., 2014).

### Cell transfection

LifeAct-mCherry and αTubulin-mCherry plasmids were kindly provided by Ana Maria Lennon. For Ecm29, a silencing siRNA kit was purchased from Qiagen (1027416) and used as the combination of 4 different siRNA at 2.5nM each one. As a control, we used a scrambled siRNA (Qiagen) at 10nM. Nucleofector R T16 (Lonza, Gaithersburg, MD) was used to electroporate 5 x 10^6^ IIA1.6 B Lymphoma cells with 2 μg of plasmid DNA. After transfection, cells were cultured for 16 hrs before functional analysis.

### Preparation of Ag-coated beads and AG-coated coverslips

Antigen-coated beads were prepared as previously described (Yuseff *et al*., 2011). Briefly, ~2x 10^7^ 3-μm latex NH_2_-beads (Polyscience, Eppelheim, Germany) were activated with 8% glutaraldehyde for four h at room temperature. Beads were washed with phosphate-buffered saline (PBS) and incubated overnight at 4°C with different ligands: using 100 μg/mL of either F(ab’)2 goat anti-mouse immunoglobulin G (IgG), referred to as BCR-Ligand^+^ or F(ab’)2 goat anti-mouse IgM, referred to as BCR-Ligand^-^ (MP Biomedical, Santa Ana, CA). For antigen extraction assays, beads were coated with BCR-Ligand^+^ or BCR-ligands^-^ plus OVA 100 μg/mL. For antigen presentation assays, beads were coated with BCR^+^ or BCR^-^ligands plus 100 μg/mL Lack protein. Antigen coverslips used to analyze the synaptic interface were coated with BCR-Ligand^+^ overnight at 4°C in PBS.

### Activation of B cells with Ag-coated beads or coverslips

Cells were plated on poly-L-Lysine– coated glass coverslips and activated with Ag-coated beads (1:1 ratio) for different time points in a cell culture incubator (37°C / 5% CO_2_) and then fixed in 4% paraformaldehyde (PFA) for 10 min at room temperature as previously described (Yuseff *et al*., 2011). Fixed cells were incubated with antibodies in PBS-0.2% BSA-0.05% Saponin. In order to measure cell spreading, the B cell line was plated onto B220/anti-IgG, or anti-IgM coated glass coverslips, respectively, for different time points at 37°C in a cell culture incubator as previously described (Reversat *et al*., 2015).

### Antigen presentation assay

Ag presentation assays were performed as previously described (Yuseff *et al*., 2011). Briefly, IIA 1.6 (I-A^d^) B cells were incubated with either Lack-BCR-Ligand^+^ or BCR-Ligand^-^ coated beads or different Lack peptide concentrations (Lack_156-173_) for 1 hr. Then Cells were washed with PBS, fixed in ice-cold PBS/0.01% glutaraldehyde for 1 min, and quenched with PBS/100mM glycine. B cells were then incubated with Lack-specific LMR 7.5 T Cells in a 1:1 ratio for 4 hrs. Supernatants were collected, and interleukin-2 cytokine production was measured using BD optiEA Mouse IL-2 ELISA set following the manufacturer’s instructions (BD Biosciences).

### Ag extraction assay

For antigen extraction assays, B cells incubated in a 1:1 ratio with BCR ligand^+^-OVA-coated beads were plated on poly-Lys cover-slides at 37°C, fixed and stained for OVA. The amount of OVA remaining on the beads was calculated by establishing a fixed area around beads in contact with cells and measuring fluorescence on three-dimensional (3D) projections obtained from the sum of each plane (Details in Image analysis section). The percentage of antigen extracted was estimated by the percentage of fluorescence intensity lost by the beads after 1 hr.

### Centrosome isolation

Centrosome from B cells was isolated as previously described (Obino *et al*., 2016) with slight modifications. Briefly, B cells in resting conditions (CLICK-2% FBS) at 37°C/5%CO_2_, were treated with adding 2μM cytochalasin D (Merck Millipore) and 0.2μM Nocodazole (Merck Millipore). Cells were washed in TBS (10mM Tris-HCl 15mM NaCl pH 7.5), then in 0.1X TBS supplemented with 8% sucrose and lysed in lysis buffer (1mM HEPES. 0.5% NP-40, 0.5 mM MgCl_2_, 0.1% β-mercaptoethanol pH 7.2) supplemented with protease inhibitors for 15 minutes. Centrosomes were isolated from post-nuclear-supernatants by consecutive centrifugations at (1) 10,000g for 30 min at 4°C on top of a 60% w/v sucrose cushion in gradient buffer (10mM PIPES, 0.1% Triton X-100, 0.1% β-mercaptoethanol pH 7.2) and (2) 40,000g for 60 min at 4°C on top of a discontinuous sucrose gradient (40-50-70% w/w). Finally, 12 fractions were recovered from the top to the bottom of the tube, and centrosome-containing fractions were detected by immunoblot γ-tubulin labeling.

### Proteasome activity

Protein extracts obtained from B cells were quantified and loaded onto black MaxiSorp 96 well plate (Nunc, Denmark) with proteasome substrate III fluorogenic (Calbiochem, Merck Millipore) diluted in Assay buffer (50mM Tris-HCl pH: 7.2, 0.05 mM EDTA, 1mM DTT). The plate was incubated for 1 hr at 37°C, and then fluorescence was measured at 360/420 nm. All measurements were performed in triplicate.

### Cell imaging

For epifluorescence imaging, all Z-stack images were obtained with 0.5 microns between slices. Images were acquired in an epifluorescence microscope (Nikon Ti2Eclipse) with an X60/1.25NA and X100/1.3NA oil immersion objectives for bead and spreading assays, respectively. For confocal microscopy, images were acquired in a Nikon Ti2Eclipse inverted microscope with 60X/1.45NA oil immersion for bead and spreading assays, with a Z-stack of 0.5 microns. For Total internal reflection fluorescence microscopy (TIRFM), images were acquired in Nikon Ti2Eclipse inverted microscope with a 100x/1.50 NA oil immersion lens and an iXON Ultra EMCCD camera at 37°C. B-cells expressing LifeAct-mCherry were plated on Ag-coated glass chambers (Nunc™ Lab-Tek™ II). Images were acquired for 30 minutes at 15 seconds per frame for spreading assay and for 1 minute at 0.75 seconds per frame for lysosome, proteasome, and actin retrograde flow tracking. For Ayriscan acquisition, images were obtained in the Zeiss Airyscan Confocal microscope with a 63X/1.4NA oil immersion lens, with a Z-stack of 0.2 μm. The images were processed using Zeiss Black Zen software and analyzed with FIJI.

### Fluorescence Recovery After Photobleaching in TIRFM (FRAP-TIRFM)

IIA1.6 cells were transfected with Actin-GFP together with scramble siRNA (siCTL) or ECM29-targeting siRNA (siECM29), and then allowed to spread onto antigen-coated coverslips for 30 to 40 min at 37°C in HEPES supplemented CLICK. Cells were then mounted on a stage-top incubator, and one central region was manually selected for photobleaching using a 405nm laser (70% intensity, 100ms, ND Stimulation unit), concurrent with Nikon TIRFM imaging. GFP signal intensity within the bleached zone was normalized to intensity values from an unbleached control region in the same cell. The curves were also y-transformed to (0,0) at t = 0 (bleaching event) so that individual recovery curves begin at the intensity minimum. Each recovery curve group was then fit to the following equation, with the constraint that Y_0_=0: Y=Y_0_+ (plateau-Y_0_)*(1-e^-k*x^). The rate constant (k) was derived by nonlinear regression analysis using GraphPad Prism software.

### Image analysis

Image processing and analysis were performed with FIJI (ImageJ) software (Schindelin *et al*., 2012), as we previously described (Ibañez-Vega, Fuentes, *et al*., 2019). The centrosome was labeled with Centrin-GFP or α-Tubulin, and determined by the brightest point where microtubules converged. Single-cell images shown in the figures were cropped from a larger field. Images brightness and contrast were manually adjusted. Centrosome polarity index was determined as previously described (Yuseff *et al*., 2011). Briefly, we manually selected the location of the centrosome (Cent) and delimited the cell and bead borders to obtain the center of mass of both CMC (Cell mass center) and BMC (Bead mass center), respectively. The position of the centrosome was projected (CentProj) on the vector defined by the CMC-BMC axis. The centrosome polarity index was calculated by dividing the distance between the CMC and CentProj and the distance between CMC-BMC. The index ranges from −1 (anti-polarized) and +1 (fully polarized).

Proteasome recruitment to the IS in bead assays was quantified by dividing the fluorescence at the bead by the whole cell’s fluorescence and then multiplying it by a factor of 100. For spreading assays, we manually delimited the border of the cell using a phalloidin label as a template (CellTemp); then, an ellipse was automatically determined (CenterTemp) at the center of CellTemp, which had a third of the CellTemp area. Next, the center’s recruitment was calculated by dividing the fluorescence normalized by its area from CenterTemp and CellTemp, subtracting 1. Therefore, positive values mean that the fluorescence is enriched at the center, and negative values, the opposite.

For actin quantification at the centrosome, we traced a circle with a 1μm radius with its center as the centrosome. The fluorescence at the centrosome (FCent) and its Area (ACent) were measured. The corresponding ratio gives the fluorescence density index (DCent=FCent/ACent). This value is divided by the density of the fluorescence of the entire cell (DCell). Values above 1 indicate an accumulation of the label at the centrosome compared to the whole cell. Whereas values below 1 indicate that there is a depletion at the centrosome compared to the whole cell.

For lysosome and proteasome tracking, we used the Trackmate plugin from FIJI (Schindelin *et al*., 2012), considering each spot with areas of 1μm^2^ and manually thresholded by the quality index.

The proteasome (Bsc2118-FL-Bodipy) fluorescence correlation with F-actin (LifeAct-mCherry) fluorescence of B cells seeded for 30 min in antigen-coated coverslips was automatically measured by a FIJI macros function. Briefly, the proteasome label was automatically detected by Analyze particle, and the proteasome and F-actin fluorescence were measured in each proteasome-spot (1μm diameter circle). Next, each fluorescence signal was normalized by the total cell fluorescence in each frame to normalize fluorescence variation by LifeAct expression or Bsc2118-FL-Bodipy dosage. Then, the proteasome fluorescence and the related LifeAct fluorescence were arranged into discrete groups and graphed.

The spreading area of LifeAct-mCherry expressing B cells activated onto antigen-coated coverslips recorded by TIRFM was assessed by FIJI. Briefly, images were thresholded and binarized to detect cell boundaries automatically, and then cell areas were detected in each frame by Analyze particle plugin (FIJI). The spreading velocity was calculated by linear regression of area per time data. The asymmetry of the immune synapse was measured by the aspect ratio of each spreading area per frame, as previously described (Kumari *et al*., 2020).

Quantification of actin retrograde flow was performed as previously described (Jankowska and Burkhardt, 2017). Briefly, TIRFM recorded LifeAct-mCherry expressing B cells after 30 min of activation onto antigen-coated coverslips, were analyzed by FIJI, by reslicing two different lamellipodia structures per each cell, and manually drawing an angle at the edge of the lamellipodium. Each angle was transformed in microns per second, by converting the angle to radians (Rad-angle) and applying the following formula: V(μm/s)=tan(Rad_angle).

Quantification of BCR clustering at the IS center was performed by using an adaptation of the clock scan analysis plugin for Fiji (Dobretsov *et al*., 2017) implemented in a personalized macro with machine learning correction with the advanced Weka segmentation tool (Arganda-Carreras *et al*., 2017). Data obtained from the images was then curated and filtered using Rstudio. Briefly, outliers were eliminated using the IQR correction and the distribution of fluorescence (MFI) was divided into 4 quartiles, considering the distance from the center of mass of each cell to their correspondent periphery. Data was assessed for its normality using the shapiro-wilk test and multiple comparison tests were performed using ANOVA and post-hoc tests (Tukey).

### Statistical analysis

Statistical analysis was performed with Prism (GraphPad Software) and RStudio. The p values were calculated using different tests, as indicated in figure legends.

## Acknowledgments

We are grateful to the Unidad de Microscopia Avanzada (UMA) of Pontificia Universidad Católica de Chile for its support in image acquisition. MI-Y was supported by a research grant from FONDECYT (**1180900**), JI-V was supported by a CONICYT PFCHA/Doctorado Nacional Chile/2015-21150566, FDV was supported by a CONICYT PFCHA/Doctorado Nacional Chile/2018-21191062.

## Author Contributions

JI-V designed, performed, and analyzed most of the experiments, assembled figures, and participated in the writing of the manuscript. FDV performed immunofluorescence and biochemical experiments and participated in the writing of the manuscript. JJ-S helped to setup image quantification protocols. JD helped with biochemical experiments. MI-Y wrote the manuscript. M-IY and AS proposed the original hypothesis, designed experiments, supervised and funded the overall research.

## Conflicts of interest

The authors declare that the research was conducted in the absence of any commercial or financial relationship that could be construed as a potential conflict of interest

## Expanded View Figure legends

**Figure EV1: Ecm29 cofractionates with the centrosome in B cells.** Representative immunoblot of centrosome fractions isolated from B cells in resting conditions, where γ-tubulin (centrosome) and Ecm29 were detected.

**Figure EV2: Ecm29 silencing does not affect levels of ubiquitylated proteins, but reduces proteasome levels at the centrosome in resting B cells. A**. Representative immunoblot of protein extracts obtained from resting B cells transfected with scrambled siRNA (siCTL) and Ecm29 targeted siRNA (siECM29). Ecm29 and BCR are shown. **B**. Quantification Ecm29 levels in A. N=7. **C**. Representative immunoblot of control and Ecm29-silenced B cells in resting conditions, stained for Ubiquitin, Ecm29, S4 (19S), αβ (20S), LMP7, and actin. **D**. Quantification of protein levels in C. N>3. **E**. Representative immunoblot of centrosome isolated fractions isolated from control and Ecm29-silenced B cells. S4 (19S) and γ-Tubulin are shown. Red rectangle indicates the centrosome-rich fractions. **F and G**. Quantification of S4 (19S) protein levels (N=3) and proteasome activity (N=2) in centrosome-rich fractions in E, respectively. **p<0.01. T-student test was performed for all statistical analyses. Mean with SEM bars are shown.

**Figure EV3: Live tracking of the proteasome shows it colocalizes with microtubules and is distributed across the IS**. **A**. Immunoblot of B cells treated with increasing concentrations of the specific proteasome probe (Bsc2118-FL-Bodipy) for 2 hours, ubiquitin and actin labeling are shown. F.C.= Fold of Change respect to the control (0nM of Bsc2118-FL-Bodipy). **B**. TIRFM Time-lapse and kymograph of α-Tubulin-mCherry (greyscale) expressing B cells probed with 5nM Bsc2118-FL-Bodipy (Red). Black arrowheads indicate proteasome positive spots. Below: Table summarizing the colocalization of the proteasome (Bsc2118-FL-Bodipy) and α-Tubulin-mCherry. Overlap coefficient, k1, k2, Manders 1, and Manders 2, are shown. **C**. Time-lapse images obtained by TIRFM of B cells labeled for proteasome (greyscale). The accumulation of proteasome tracks (right) is shown. The coldest colors represent the fastest tracks—Kymograph (below). Black arrowheads indicate proteasome positive spots at center and periphery. **D**. Histogram of proteasome diffusion coefficient measured in C. N=2, Cells>30. Scale Bar= 10μm.

**Figure EV4: Proteasome recruitment and distribution at the IS rely on Ecm29: A.** Representative Time-lapse images by TIRFM of control (siCTL) and Ecm29-silenced (siECM29) B cells after 30 min of activation on antigen-coated coverslips, probed with 5nM Bsc2118-FL-Bodipy (grey scale). The cell boundary (black line) delimited by the LifeAct-mCherry signal (mask) and the tracks’ accumulation are shown. The coldest colors represent the fastest tracks. **B**, **C**, **D, and E**. Quantification of track displacement, track duration, track mean velocity, and the number of tracks of proteasomes at the IS of control or Ecm29 silenced B cells after 30 min of activation on antigen-coated coverslips. N>20 Cells. *p<0.05, **p<0.01, ****p<0.0001. Unpaired T-Test was performed. Scale Bar = 10μm.

**Figure EV5: Proteasome arrival at the IS negatively regulates actin polymerization. A**, **C, E, and G.** Representative TIRFM images of control, MG-132 pre-treated (5μM for 1 hour), Bsc2118-FL-Bodipy overdosed (100nM), and Ecm29-silenced (siECM29) B cells. B cells were plated for 30 min onto antigen-coated coverslips and then recorded. LifeAct-mCherry (white) and proteasome (green [Bsc2118-FL-Bodipy]), magnifications proteasome spots (white dashed rectangles), together with their respective kymograph, are shown. White arrowheads indicate the proteasome arrival at the IS close to F-actin structures. **B**, **D, F, and H**. Quantification of the proteasome (green) and LifeAct-mCherry (Red) distribution at proteasome positive spots (white dashed line) and their distribution in time (Kymograph) showed on A, C, D, and G, respectively. **I**. Quantification of normalized fluorescence intensity correlation between Bsc2118-FL-Bodipy and LifeAct-mCherry, on Bsc2118-FL-Bodipy-positive spots of 1μm in diameter shown in A, C, G, and E. N>10. *p<0.05. **p<0.001. ***p<0.0005. ****p<0.0001. 2-ways ANOVA with Sidak’s post-test was performed. Each dot represents an independent positive Bsc2118-FL-Bodipy spot. Mean with SEM lines are shown. Scale Bar = 10μm.

**Figure EV6: Centrosome-nucleus separation and centrosome recruitment to the IS is impaired in Ecm29-silenced B cells: A**. Representative Epifluorescence images of control (siCTL) and Ecm29-silenced (siECM29) B cells, activated with antigen-coated beads for different time points. Centrosome (Centrin-GFP) and nucleus (DAPI) are shown. The dashed line represents the distance between the centrosome-nucleus. **B and C**. Quantification of centrosome-nucleus distances and centrosome polarization in A. N>3. Cells>116. Each dot represents an individual measurement (C). Mean with SEM lines (C) and bars (B) are shown. ***p<0.0005. ****p<0.0001. Two-way ANOVA with Sidak’s post-test. Scale Bar= 10μm.

